# Strand asymmetry of DNA damage tolerance mechanisms

**DOI:** 10.1101/2024.01.21.576515

**Authors:** Juan Carlos Cañas, Dolores Jurado-Santiago, Mohammed Al Mamun, María P. Sacristán, Esther C. Morafraile, Javier Zamarreño, Katsunori Fujiki, Katsuhiko Shirahige, Avelino Bueno, Rodrigo Bermejo

## Abstract

DNA damage tolerance mechanisms are crucial for timely and accurate chromosomal replication in response to DNA polymerase stalling. Ubiquitylation of the replicative sliding clamp PCNA drives major tolerance pathways, error-free homologous recombination template switching and error-prone translesion synthesis, though their dynamics at forks and pathway choice determinants are poorly understood. Using strand-specific genomics we revealed an asymmetric nature of tolerance pathways, characterized by preferential template switching-driven recombinase engagement of stalled nascent lagging strands and translesion synthesis usage in response to leading strand polymerase stalling. This asymmetry, determined by a strand-dynamic interplay between PCNA-ubiquitin writers and erasers, likely stems from necessities dictated by leading and lagging strand replication mechanisms and has implications for asymmetric mutation inheritance.

**One-Sentence Summary:** DNA damage tolerance mechanisms respond asymmetrically to leading or lagging strand polymerase blocks.

## Main Text

DNA damage tolerance (DDT), or post-replicative repair, mechanisms allow cells to efficiently overcome blocks to replicative polymerase synthesis, crucial for timely chromosome duplication and integrity maintenance **(*1-3*)**. Chromosome replication is primary carried out by DNA polymerase ε (Pol ε), which continuously synthetizes leading nascent strands, coupled to CMG (Cdc45-MCM-GINS) helicase unwinding of parental duplexes, and DNA polymerase δ (Pol δ), which extends short primers generated by DNA polymerase α/primase to form Okazaki fragments, later matured into continuous lagging nascent strands **(*4, 5*)**. Endogenous and exogenous factors causing base lesions, deoxy-nucleotide (dNTP) shortage or secondary DNA structure stabilization can stall synthesis by replicative polymerases, prompting DDT pathways. Central to tolerance mechanisms are post-translational modifications of the DNA sliding clamp homotrimer PCNA (*Proliferating Cell Nuclear Antigen*) (Fig. S1A)**(*6*)**. Uncoupling between DNA unwinding and synthesis following polymerase stalling generates extended ssDNA tracks **(*7*)**, which upon coating by RPA (Replication Protein A) recruit the Rad18 E3-ubiquitin ligase **(*8*)**. Rad18, in complex with the E2-ubiquitin conjugase Rad6, attaches single ubiquitin moieties to lysine164 of PCNA **(*9*)**. K164-monoUbiquitylated PCNA recruits translesion synthesis (TLS) polymerases, Rev1, Pol η (Rad30), and Pol ζ (Rev3, Rev7, Pol31, and Pol32) in budding yeast, which act sequentially to mediate bypass of diverse blocks **(*10*)**. Owing to TLS polymerase low fidelity and lack proofreading activity, Translesion Synthesis tolerance is largely error-prone **(*11*)**. Extension of K164-monoUbi-PCNA to K63-linked polyubiquitin chains is mediated by Ubc13-Mms2 E2-conjugase and the E3-ligase Rad5 **(*9*)**. K164-polyubiquitylated PCNA inhibits Translesion Synthesis and promotes block circumvention using the sister chromatid as a template for synthesis **(*12, 13*)**. This Rad51 and Rad52 recombinase-dependent process is known as Template Switching (TS) and allows error-free bypass of polymerase blocks. In addition, Rad5 promotes TLS, independently of its ubiquitin ligase activity, by stabilizing TLS polymerases at block sites **(*14-16*)**. Even if better studied in the context of DNA lesion bypass, DDT pathways are also crucial to promote cell viability and genome integrity in response to “lesion-less” polymerase blocks **(*17*)**.

Different PCNA-interacting factors hold a potential to downregulate DDT mechanisms. Fork-associated ubiquitin proteases Ubp10 and Ubp12 diversely reverse PCNA ubiquitylation **(*18, 19*)**. Ubp10 fully removes ubiquitin chains form lysine 164, while Ubp12 trims K63-linked ubiquitin, accumulating monoubiquitylated-PCNA **(*20*)**. The alternative clamp loader Replication Factor C-like complex Elg1-RFC unloads PCNA in response to polymerase stalling, a process that operates preferentially on lagging strands **(*21, 22*)**. Lastly, the Ubc9 SUMO-conjugating enzyme and the Siz1 E3-SUMO ligase attach small ubiquitin-like SUMO molecules to PCNA lysine164 and lysine 127 during unperturbed replication **(*9*)**. SUMOylated PCNA recruits the anti-recombinogenic helicase Srs2, which limits salvage recombination at replication forks by disrupting Rad51 recombinase-nucleofilaments and inhibiting Rad52 **(*23, 24*)**.

While key actors and pathways have been genetically characterized, DDT dynamics and mechanism-choice determinants at blocked polymerase sites remain poorly understood. Hence, we studied the functional architecture of tolerance mechanisms elicited by replicative polymerase stalling using strand-specific genomics. We report an asymmetric nature of the DDT response, characterized by preferential usage of homologous recombination-based Template Switching in response to lagging strand synthesis stalling and TLS polymerase recruitment to leading strand polymerase blocks.

## Results

### Strand-asymmetric DNA damage tolerance response to replicative polymerase blocks

We exploited eSPAN (*enrichment and Sequencing of Protein-Associated Nascent Strands*) (Fig. S1B), which allows differentially determining differential protein binding to nascent DNA strands **(*25*)**, to examine DDT effector recruitment to sites of replicative polymerase stalling in yeast cells. For eSPAN, bromodeoxyuridine (BrdU)-labelled nascent chromatin is crosslinked and sheared, prior to immunoprecipitation with antibodies against a protein of interest (ChIP). Newly replicated DNA is then isolated through BrdU-immunoprecipitation (BrdU-IP) from input and ChIP DNA to obtain BrdU-IP and eSPAN fractions containing, respectively, all nascent strands or nascent strands associated to the protein of interest. Strand-specific sequencing determines the abundance of Watson and Crick reads, unambiguously assigned to leading or lagging nascent strands in BrdU-IP and eSPAN fractions based on positional alignment in respect to the polarity of DNA synthesis away from origins. Pilot eSPAN experiments targeting the catalytic subunits of Polε (Pol2) or Polδ (Pol3) evidenced their asymmetric association to stalled nascent strands (Fig. S1C and S1D) **(*25*)**, in agreement with the respective primary roles of these enzymes in leading and lagging strand replication **(*26, 27*)**.

To asses stalled nascent strand engagement in Homologous Recombination-mediated DNA damage tolerance events, we first performed eSPAN using antibodies specific to the Rad51 recombinase **(*28*)**. Upon induction of polymerase stress by cell release into S-phase in the presence of the dNTP pool-depleting drug hydroxyurea (HU), Rad51-ChIP reads clustered at positions of DNA synthesis stalling around active replication origins, as evidenced by BrdU-IP read mapping (Fig. 1A). Indicative of recombinase engagement of stalled nascent strands, Rad51-eSPAN sequencing reads also sharply clustered around replication origins. Averaged Rad51-ChIP and BrdU-IP read enrichments overlapped genome-wide, while a higher abundance for Watson-left and Crick-right (lagging strand) reads was detected in eSPAN profiles (Fig. 1B). Accordingly, Watson over Crick read ratios rendered positive and negative values, respectively, leftwards and rightwards to origins (Fig. 1C), of a magnitude equivalent to those obtained for Polδ (Fig. S1D), and a significantly higher abundance of lagging-strand reads was detected at forks emanated from a vast majority of origins (Fig. 1D). These data reveal a marked preference for recombinase engagement in response to lagging-strand polymerase stalling. Rad51 preferential lagging-strand association was also observed when polymerase-blocking lesions were induced by treatment with the alkylating agent methyl-methanesulphonate (MMS) (Fig. S2A), despite reduced fork synchronicity dispersing Rad51-ChIP reads. Hence, our results indicate a preference for homologous recombination-based mechanism usage in response to lagging strand synthesis blocks.

**Fig. 1.**
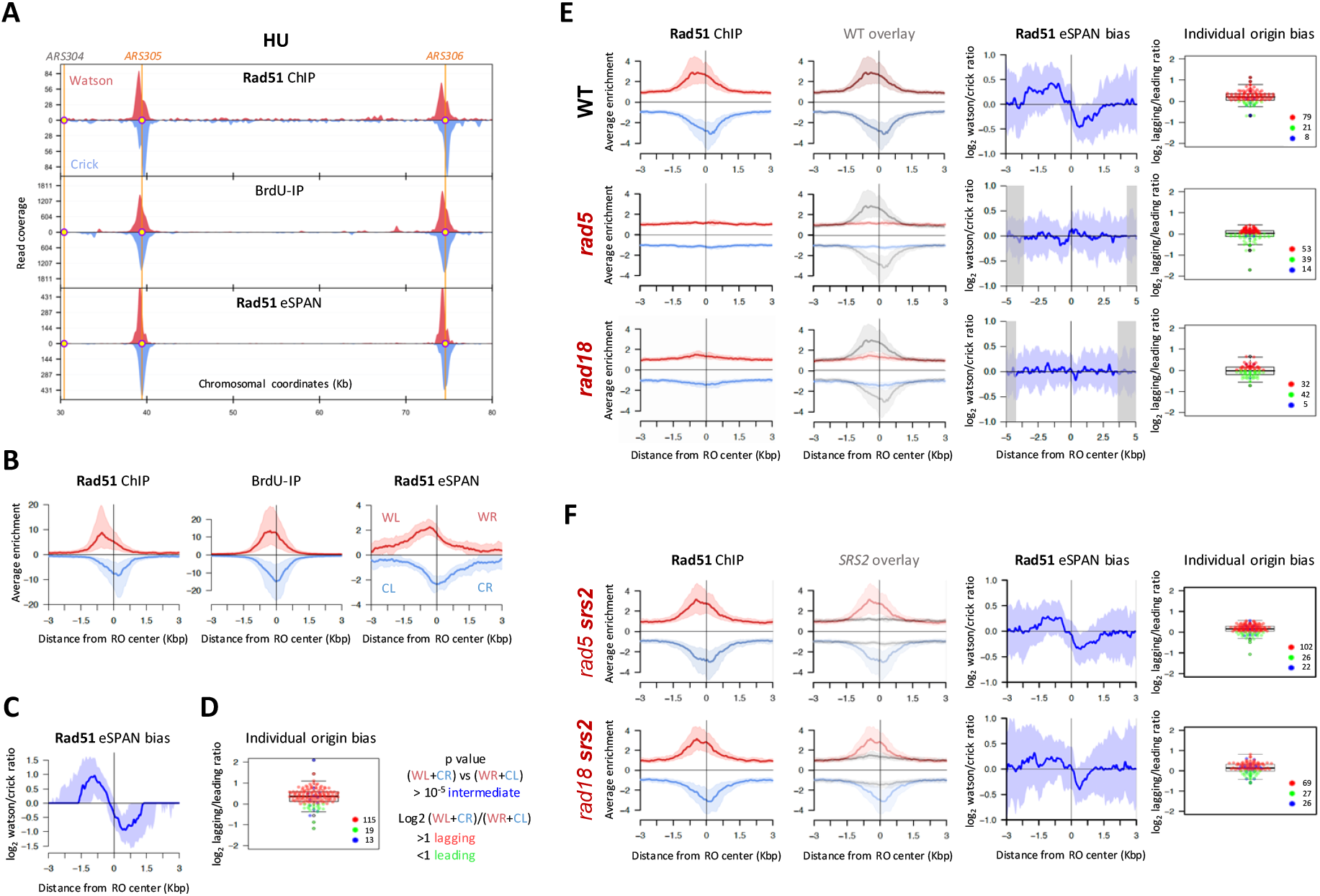
Homologous Recombination-mediated DNA damage tolerance mechanisms preferentially engage stalled lagging strands. (**A-D**) Rad51-eSPAN analysis in cells collected one hour after release from G1 in the presence of 0.2M HU. (**A**) Rad51-ChIP, BrdU-IP and Rad51-eSPAN fraction read coverage plotted along a region spanning from 30 to 80 Kb on chromosome III. Positions of early replication origins *ARS305* and *ARS306* (orange), as well as the dormant origin *ARS304* (grey) are indicated. Watson and Crick reads are plotted in red and blue, respectively. (**B**) Averaged input-normalized Rad51-ChIP and BrdU-IP, as well as BrdU-IP-normalized Rad51-eSPAN read enrichment ratios around replication initiation sites. Median (red-blue) and 25-75 percentile (shaded) enrichment values plotted around BrdU-IP peak summits. (**C**) Averaged Rad51-eSPAN strand read ratios around replication initiation sites. Median (deep blue) and 25-75 percentile (shaded) BrdU-IP-normalized Watson over Crick read log2 ratio values plotted around BrdU-IP peak summits. (**D**) Individual replication origin leading-strand or lagging-strand biases. Dot and box plots showing the variation of BrdU-IP normalized lagging over leading eSPAN read ratios around BrdU-IP peak summits. Each dot corresponds to an individual origin-containing eSPAN peak, falling into one of three categories (intermediate, lagging-strand bias or leading-strand bias) based on lagging over leading read binomial distribution significance and bias calculation (as shown). (**E**) eSPAN analysis of Rad51 association to replicative polymerase stalling sites in WT, *rad5Δ* and *rad18Δ* cells as in A-D. Averaged input-normalized Rad51-ChIP read enrichment profiles, alone or overlayed to WT profiles (in grey), as well as averaged Rad51-eSPAN strand read ratios and individual replication origin leading-strand or lagging-strand biases are shown. (**F**) eSPAN analysis of Rad51 association to replicative polymerase stalling sites in *rad5Δ srs2Δ* and *rad18Δ srs2Δ* cells. Averaged input-normalized Rad51-ChIP read enrichment profiles, alone or overlayed to *rad5Δ* / *rad18Δ* (*SRS2*) profiles (in grey), as well as averaged Rad51-eSPAN strand read ratios and individual replication origin leading-strand or lagging-strand biases are shown

We next performed Rad51 eSPAN in *RAD5*- and *RAD18*-ablated cells, respectively defective in either PCNA poly- or mono- and poly-ubiquitylation. Absence of either K164-ubiquitin writer resulted in a drastic reduction of Rad51enrichment at sites of stalled synthesis (Fig. 1E), precluding preferential strand association assessment. Quantitatively, recombinase association to sites of polymerase stalling was reduced to background levels in PCNA-K164 ubiquitylation-deficient *rad18* and *rad5* mutants (Fig. S2B). Hence, recombinase engagement of lagging nascent strands in response to polymerase blockage mostly corresponds to Template Switching events. We exploited the lack of Rad51 stalled synthesis site binding in ubiquitin-ligase mutants to also gain insight on nascent strand engagement by the salvage recombination pathway. We performed Rad51-eSPAN in cells lacking TS, owing to *rad5* or *rad18* deletion, in combination with ablation of the Srs2 helicase. While the absence of Srs2 alone did not lead to major changes in Rad51 enrichment or nascent strand association preference (Fig. S2C), *rad5* and *rad18* mutants exhibited a marked recovery of recombinase enrichment at polymerase stalling sites (Fig. 1F). Quantitatively, Srs2 ablation salvaged approximately one third of wild-type Rad51 association in *rad5* and *rad18* cells, while it did not result in a significant increase in PCNA-K164 ubiquitin writer proficient cells (Fig. S2D), possibly reflecting a reduced proficiency of salvage mechanisms in responding to lagging-strand polymerase blocks compared to Template Switching. Of note, Rad51 also exhibited preferential association to lagging nascent strands in salvage pathway-permissive *rad5 srs2* and *rad18 srs2* genetic contexts, pointing at a preference for Homologous Recombination-based DDT usage to overcome lagging strand polymerase blocks.

We next analyzed Translesion Synthesis strand distribution by carrying out eSPAN experiments targeting the Rev1 TLS polymerase and TLS polymerase scaffold. In consistence with previous reports **(*14*)**, Rev1-ChIP read coverage also clustered around active replication origins, overlapping nascent strand synthesis stalling positions, as determined by BrdU-IP read mapping (Fig. 2A). Normalized Rev1 ChIP enrichment spanned sites of replicative synthesis stalling around active replication origins genome-wide (Fig. 2B), and a higher abundance for Crick-left and Watson-right reads was observed in Rev1-eSPAN read enrichment profiles, revealing a preferential recruitment of Rev1 to sites of leading strand polymerase stalling. In agreement, Watson over Crick eSPAN read ratios rendered negative and positive values, respectively, leftwards and rightwards to fired origins, and a significantly higher abundance of leading-strand reads was detected at forks emanated from the majority of individual replication origins (Fig. 2C). eSPAN analysis of TLS polymerase η (Rad30), also evidenced enrichment at sites of nascent strand synthesis stalling upon HU treatment, although it rendered an overall non-significant leading bias trend (Fig. S3A). We note that a nascent strand association bias could not be determined for Rev1 in cells replicating in presence of MMS, likely owing to fork dispersion, which precluded its detection at sites of stalled DNA synthesis (Fig. S3B). Next, we tested TLS-pathway genetic dependencies for Rev1 recruitment. Ablation of the Rad18 ubiquitin ligase, impairing PCNA-K164 mono-ubiquitylation, largely abolished Rev1enrichment around active replication origins (Fig. 2D and S3C). In turn, deletion of *RAD5* resulted in a marked but not complete reduction of Rev1 enrichment at sites of DNA synthesis stalling (Fig. 2D), with 10-30% of wild type Rev1 association retained in *rad5* cells as compared to *rad18* mutants (Fig. S3C), likely reflecting the fraction of Rev1 engagement dependent on Rad5 for recruitment **(*14*)**. Residual Rev1 recruitment rendered a slight non-significant trend for leading strand association in *rad5* cells, hinting at a strand preferential usage for Rad5-independent TLS events.

**Fig. 2.**
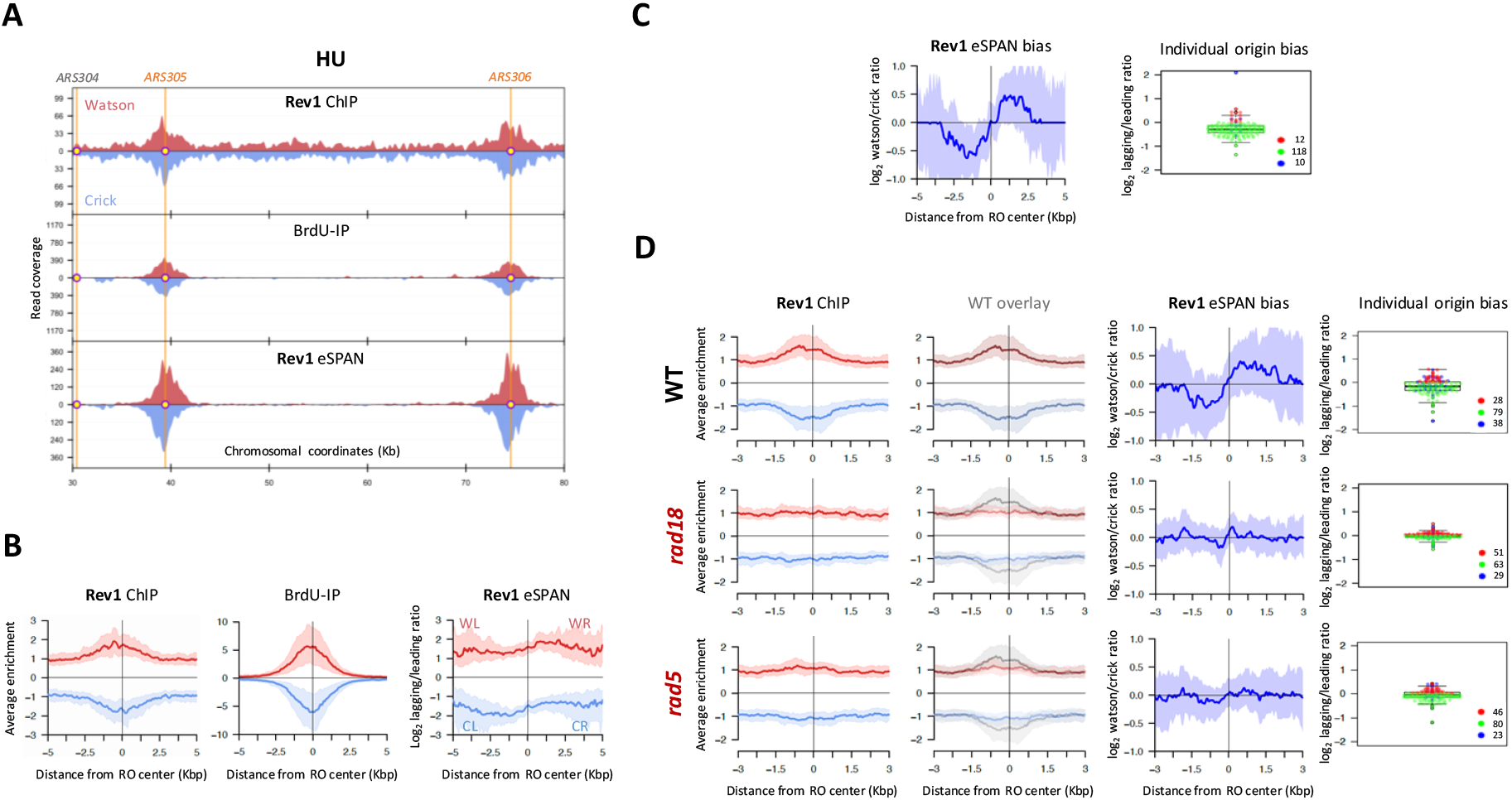
Translesion Synthesis polymerase preferential recruitment to leading strand polymerase block sites. (**A-C**) eSPAN analysis of Rev1 association to replicative polymerase stalling sites. Rev1-Myc cells were released from G1 in the presence of 0.2M HU and collected after one hour. (**A**) Rev1-ChIP, BrdU-IP and Rev1-eSPAN fraction read coverage plotted along a region spanning from 30 to 80 Kb on chromosome III. (**B**) Averaged input-normalized Rev1-ChIP and BrdU-IP, as well as BrdU-IP-normalized Rev1-eSPAN read enrichment ratios around replication initiation sites. (**C**) Averaged Rev1-eSPAN strand read ratios around replication initiation sites and individual replication origin leading-strand or lagging-strand biases. (**D**) eSPAN analysis of Rev1 association to replicative polymerase stalling sites in Rev1-Flag WT, *rad18Δ* and *rad5Δ* cells as in A-C. Averaged input-normalized Rev1-ChIP read enrichment profiles, alone or overlayed to WT profiles, as well as averaged Rev1-eSPAN strand read ratios and individual replication origin leading-strand or lagging-strand biases are shown.

These results evidence TLS polymerase usage for the on-the-fly response to leading strand polymerase blocks and, taken together with TS preferential engagement of stalled lagging strands, demonstrate a strand-asymmetric nature of the DNA damage tolerance response.

### Dynamic interplay between PCNA-K164 ubiquitin writers and erasers in DDT asymmetry determination

We next investigated the determinants of strand-asymmetric DDT pathway choice. Given the central role of PCNA ubiquitylation in driving tolerance mechanisms, we first investigated PCNA and K164 ubiquitin-writer association to stalled nascent strands in a dynamic framework. We set up conditions to monitor PCNA unloading from chromatin upon polymerase stalling by carrying out Pol30-eSPAN 40 and 60 minutes after release into a synchronous S-phase in the presence of HU (Fig. 3A and Fig. S4A). At an earlier stage (40’), PCNA enriched at sites of polymerase stalling around replication origins with a neutral bias and a slight overall lagging strand-preferential association trend, resembling what observed in unloading-deficient *elg1* mutants **(*25*)**. Twenty minutes later (60’), Pol30-eSPAN bias shifted toward a net preference for leading strand association, indicating that PCNA unloading from stalled lagging strands takes place during this time interval. We next examined Rad18 and Rad5 ubiquitin-writer association to sites of polymerase stalling in the frame of PCNA unloading. Quantitative-ChIP evidenced maximum PCNA-ubiquitin writer recruitment levels 40’ after release into an HU-challenged S phase, followed by a gradual decline coincident with PCNA unloading timings (Fig. S4B).

**Fig. 3.**
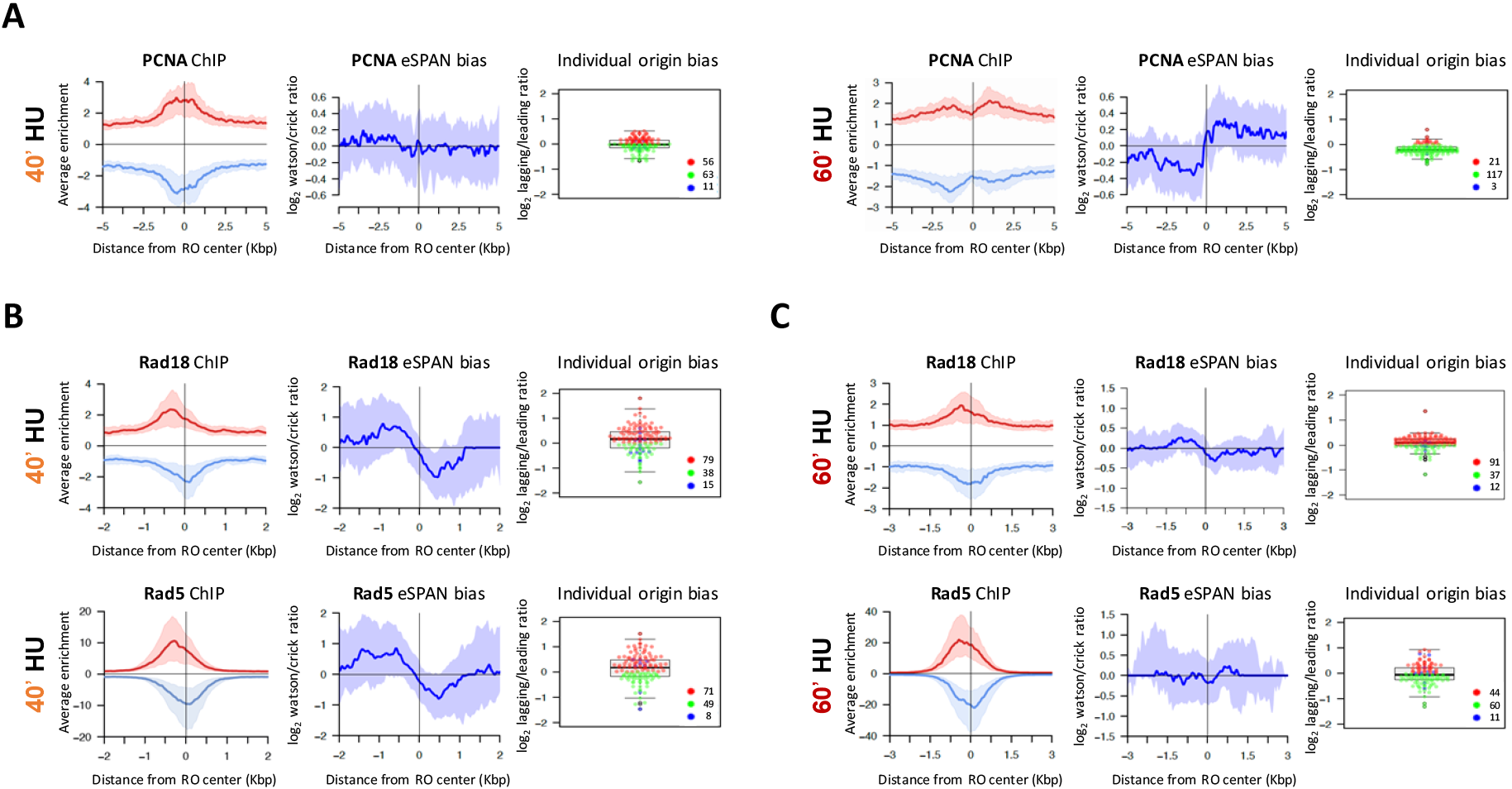
Dynamic association of PCNA and PCNA-K164 ubiquitin writers to stalled nascent strands. (**A**) eSPAN analysis of PCNA association to replicative polymerase stalling sites. Pol30-Flag cells were released from G1 in the presence of 0.2M HU and collected after 40 or 60 minutes. PCNA-ChIP read enrichment profiles, as well as averaged PCNA-eSPAN strand read ratios and individual replication origin leading-strand or lagging-strand biases are shown. (**B-C**) eSPAN analysis of Rad18 and Rad5 association to replicative polymerase stalling sites. Rad18-Flag or Rad5-Flag cells were released from G1 in the presence of 0.2M HU and collected after 40 (**B**) or 60 (**C**) minutes. Rad18/Rad5-ChIP read enrichment profiles, as well as averaged Rad18/Rad5-eSPAN strand read ratios and individual replication origin leading-strand or lagging-strand biases are shown.

Accordingly, eSPAN analysis evidenced a genome-wide recruitment of Rad18 and Rad5 ligases to sites of polymerase stalling prior to PCNA unloading (40’), both characterized by a strong bias toward lagging nascent strand association (Fig. 3B). Simultaneous ubiquitin writer recruitment is likely to generate a PCNA polyubiquitylation-promoting environment stimulating lagging strand engagement in TS events. In agreement with this notion, Rad51 enriched at sites of replication stalling at this early stage with a stark preference for lagging strand association (Fig. S4C). Upon PCNA unloading, preferential association to stalled nascent strands of the ligases shifted, resulting in a slight lagging bias for Rad18 and a neutral average bias for Rad5 (Fig. 3C). Together, overall association level reduction (Fig. S4B) and shift toward a more symmetric binding to nascent strands suggest that PCNA-ubiquitin writers dissociate from stalled lagging strands along with PCNA. Rad18 dissociation is not likely due to a relative decrease in ssDNA at lagging strands, as RPA subunit Rfa1 preferential bias was unchanged during this time (Fig. S4D). Taken together, this evidence suggests that simultaneous recruitment of Rad18 and Rad5 ubiquitin ligases enables lagging nascent strand engagement in TS recombination by promoting PCNA polyubiquitylation at sites of polymerase stalling. In addition, it raises the possibility that PCNA unloading serves as a mechanism precluding further enrolment of DDT effectors to TS-engaged lagging strands.

The fact that Rad18 and Rad5 exhibit neutral eSPAN biases following PCNA unloading implies that they are also recruited to stalled leading nascent strands. Nonetheless, Rad51 retains a stark preference for lagging strand association at later stages of polymerase stalling (Fig. 1), suggesting the existence of mechanisms counteracting recombinase engagement at leading strands. It was proposed that Ubp10 and Ubp12 PCNA-K164-Ubiquitin erasers downregulate DDT events at stalled forks **(*20*)**. Hence, we performed eSPAN experiments targeting Ubp10 and Ubp12 de-ubiquitylases (DUBs) in the frame of PCNA unloading. At early stages, both DUBs enriched at sites of replicative polymerase stalling and exhibited slight lagging strand preferential association biases (Fig. 4A), reminiscent of that observed for PCNA (Fig. 3A). Since Rad51 preferentially associates lagging strands at this stage, Rad18 and Rad5 Ubi-writers most likely prevail over erasers, resulting in PCNA poly-ubiquitylation at lagging strand polymerase stalling sites. Following PCNA unloading, Ubp10 showed a neutral eSPAN bias, implying an equivalent association of the eraser to leading and lagging nascent strands. In contrast, by 60’ Ubp12 exhibited a slight preference for leading strands (Fig. 4B). These data suggest that PCNA-K164 erasers also dissociate from lagging nascent strands along with PCNA, with a comparatively higher retention of Ubp10, perhaps related to physical interaction with lagging strands synthesis machineries **(*29*)**. Furthermore, this evidence argues for the existence of an eraser-differential environment at stalled leading strands that, based on Ubp12 preference to trim polyubiquitin chains **(*20*)**, may favor mono-ubiquitylated PCNA accumulation and, hence, TLS polymerase recruitment. Since both Ubp10 and Ubp12 oppose PCNA poly-ubiquitylation, we tested whether these DUBs cooperate to limit Rad51 recruitment to stalled leading nascent strands by carrying out Rad51 eSPAN in *ubp10 ubp12* double deletion mutants (Fig. 4C). In contrast to wild type cells, in absence of both erasers Rad51 displayed a symmetrical nascent strand association bias and an increase in the proportion of origins showing a preferential leading association. This indicates that Ubp10 and Ubp12 counteract TS usage in response to leading strand blocks, likely by promoting PCNA de-ubiquitylation. Taken together, these data evidence a dynamic strand-differential interplay between PCNA-K164 ubiquitin writers and erasers determining a starkly asymmetric DDT response to replicative polymerase blocks (Fig. 4D). On the one hand, Rad18 and Rad5 are predominantly attracted to stalled lagging strands prior to PCNA unloading, to cooperatively promote PCNA polyubiquitylation and engagement in template switching. On the other, Ubp10 and Ubp12 favor TLS usage over TS engagement, likely by downregulating steady-state leading strand poly-ubiquitylated PCNA levels and, hence, counteracting engagement by Rad51.

**Fig. 4.**
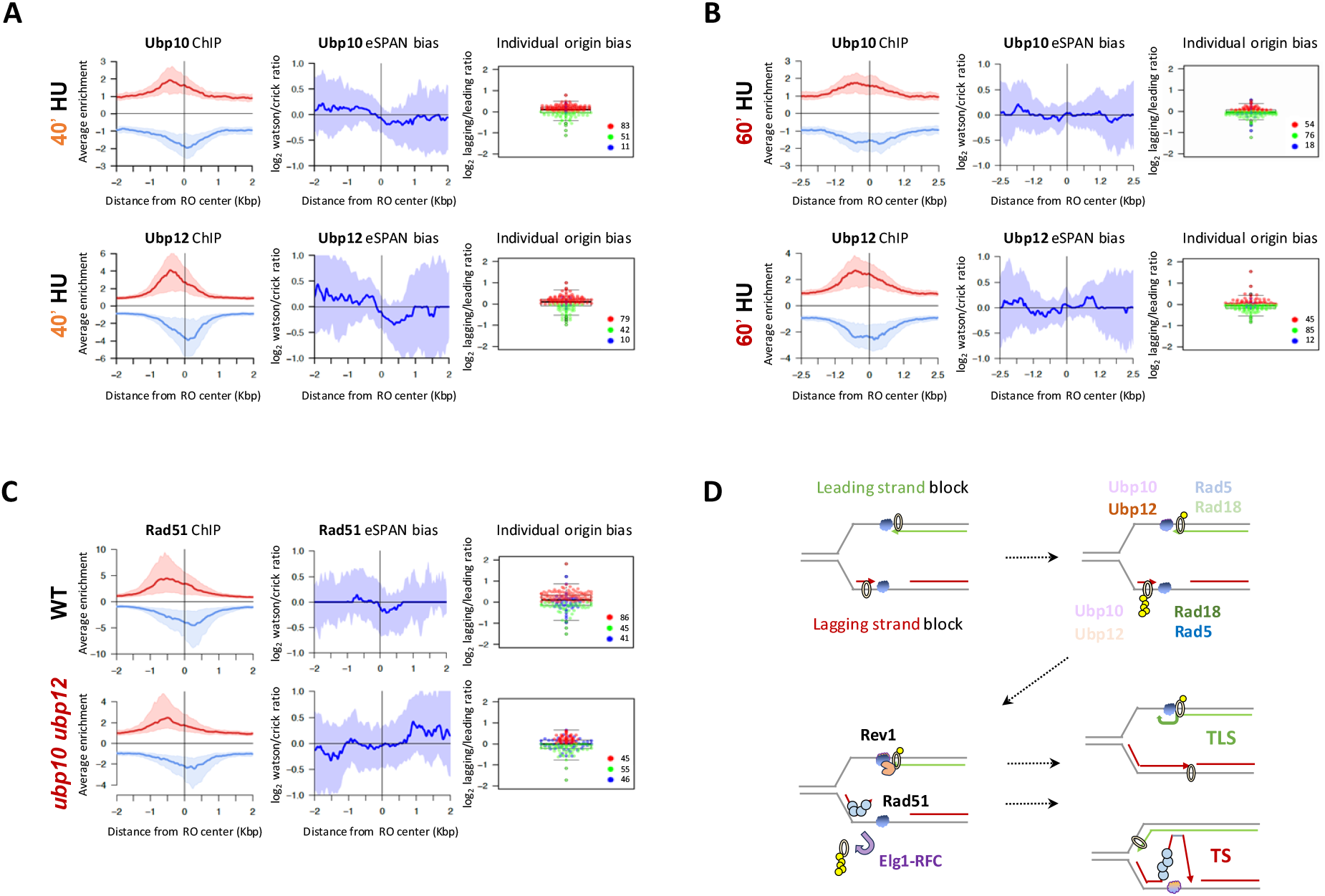
PCNA-K164 de-ubiquitylases Ubp10 and Ubp12 promote preferential lagging nascent strand engagement by Rad51. (**A-B**) eSPAN analysis of Ubp10 and Ubp12 association to replicative polymerase stalling sites. Ubp10-Myc and Ubp12-Myc cells were released from G1 in the presence of 0.2M HU and collected after 40 (**A**) or 60 (**B**) minutes. Ubp10/Ubp12-ChIP read enrichment profiles, as well as averaged Ubp10/Ubp12-eSPAN strand read ratios and individual replication origin leading-strand or lagging-strand biases are shown. (**C**) eSPAN analysis of Rad51 association to replicative polymerase stalling sites in WT and *ubp10Δ ubp12Δ* cells. Rad51-ChIP read enrichment profiles, as well as averaged Rad51-eSPAN strand read ratios and individual replication origin leading-strand or lagging-strand biases are shown. (**D**) Working model for the interplay of PCNA, PCNA-K164 ubiquitin writers and erasers in determining DNA damage tolerance mechanism strand asymmetry in response to replicative polymerase blocks (see text for details).

Here we demonstrate an asymmetric behavior of DDT mechanisms in response to polymerase blocks, characterized by a higher engagement of stalled nascent lagging strands by the Rad51 recombinase that follows Template Switching genetic requirements. This preference is observed upon induction of HU-induced dNTP shortage or base-alkylation MMS treatment, in which both leading and lagging strand polymerases should be similarly affected. Hence, Template Switching appears to be a major pathway responding to events blocking DNA polymerase δ during lagging strand replication. Rad51 engagement of lagging strands occurs prior to, and persists after, PCNA unloading, suggesting an early commitment of these strands in HR-driven Template Switching, which is likely a consequence of concomitant Rad18 and Rad5 association to lagging strands, generative of a PCNA-polyubiquitylation promoting environment. Since PCNA-monoubiquitylation is a key event stimulating TLS polymerase recruitment, unloading from lagging strands by the Elg1-RFC alternative complex likely precludes further engagement in TLS. Accordingly, Rev1 polymerase and TLS scaffold shows a preference for stalled leading strand association, suggesting a preponderant TLS usage to cope with DNA polymerase ε stalling. However, we note that the magnitude of Rev1 TLS polymerase bias is relatively smaller compared to that observed for Rad51 and that our eSPAN data do not rule out some extent of TLS usage in response to lagging strand blocks as observed in yeast *in vitro* reconstituted systems **(*30*)**.

In our experimental setup, DDT effector recruitment to polymerase blocks is detected genome-wide, typically at around 300 fork stalling sites flanking active replication origins, corresponding to an approximate total of about 600 blocked leading and lagging strand polymerases. Hence, these polymerase block-induced events diverge from previously reported post-replicative repair of ssDNA gaps, which cluster in orders of magnitude smaller (1-3) number of foci and are uncoupled from sites of DNA synthesis **(*31*)**, and likely reflect on-the-fly tolerance attempts. We propose that DDT usage asymmetry stems from the semi-discontinuous nature of chromosomal DNA replication. TS requires the invasion of the sister chromatid to use it as a template for error-free synthesis opposite to the block **(*1, 3*)**. Hence, TS usage for on-the-fly bypass of leading strands polymerase blocks might be risky, as the lagging strand serving as a synthesis template is likely to present discontinuities owing to ssDNA gaps or un-ligated nicks between Okazaki fragments. Consequently, leading strand TS attempts could result in premature cessation of bypass DNA synthesis or lead to chromosomal aberrations, owing to polymerase slippage or aberrant recombination cycles, and would be therefore deterred. Conversely, preferential TLS usage in response to leading strand blocks might reflect an advantage in maintaining DNA synthesis coupled to replicative helicase unwinding. In this respect, we note that repriming of leading strand synthesis is very inefficient in *in vitro* reconstituted systems **(*32*)**. Leading strand priming is likely to also be inefficient *in vivo*, as current models for leading strand synthesis initiation imply origin-proximal Okazaki fragment extension by DNA polymerase δ, which is handed over to CMG-associated DNA polymerase ε complex without intervention of the Polα/Primase complex **(*33*)**. In this view, leading strand synthesis continuity and helicase coupling might be favored by on-the-fly TLS polymerase bypass and swift replicative polymerase handover **(*32*)**. At lagging strands, intrinsic uncoupling from unwinding by CMG would reduce this advantage of TLS usage, while providing sufficient time to engage in error-free TS events and counteract mutagenesis. In metazoan cells, efficient repriming can also be carried out by PrimPol, absent in yeast **(*34*)**. It remains to be determined whether PrimPol-mediated re-priming can occur on leading strands, and whether, hence, preferential TLS usage is maintained in response to leading strand polymerase blocks **(*35*)**. In particular, considering that the TS/TLS strand-differential usage might lead to an asymmetric mutagenic load inheritance, with potential implications in contexts such as organismal development or tumor proliferation.

## Acknowledgments

We thank P. Sung, J.A. Tercero and A. Serra for reagents, strains and technical advice, and all members of our laboratories for insightful discussions.

## Funding

Spanish Ministry of Science grant PID2020-116003GB-100 (RB)

Spanish Ministry of Science grant PID2019-109616GB-100 (AB, MPS)

*Junta de Castilla y León* SA042P17 / SA103P20 (AB, RB)

*Junta de Castilla y León* Postdoctoral Fellowship (JCC)

Spanish Ministry of Science *Formación del Personal Investigador* PRE2018-084025 (DJS)

Spanish Ministry of Science *Juan de la Cierva - Incorporación* Postdoctoral Fellowship IJC2019-041728-I (ECM)

*Junta de Castilla y León* Predoctoral Fellowship (JZ)

Swedish Research Council grant 2022-03478 (KS)

MEXT grant 20H05940 (KS)

AMED BINDS grant 22ama121020j0001 (KS)

## Author contributions

Conceptualization: RB, AB, MPS, JCC, MAM

Data curation: MAM, DJS, RB

Formal Analysis: MAM

Investigation: JCC, DJS, MPS, ECM, JZ, KF

Methodology: RB, DJS, MAM

Project administration: RB, AB, MPS, KS

Software: MAM, RB

Supervision: RB, AB, MPS, KS

Visualization: JCC, RB

Funding acquisition: RB, AB, KS

Writing – original draft: RB

Writing – review & editing: RB, AB, JCC, DJS, MAM, MPS, ECM, JZ

## Competing interests

Authors declare that they have no competing interests.

## Data and materials availability

Deep sequencing datasets have been deposited in the Gene Expression Omnibus (GEO) (GSE253317). Code used in the analysis was deposited in GitHub **(*47*)**.

## Supplementary Materials

Materials and Methods

Tables S1 to S4

Figs. S1 to S4

## Notes

https://www.ncbi.nlm.nih.gov/geo/query/acc.cgi?acc=GSE253317

https://github.com/MohammedAlMamun/ngsAnalyser1.1.4

### Competing Interest Statement

The authors have declared no competing interest.

## References

1. D. Branzei, I. Psakhye, DNA damage tolerance. Curr Opin Cell Biol 40, 137–144 (2016).

2. N. Garcia-Rodriguez, R. P. Wong, H. D. Ulrich, Functions of Ubiquitin and SUMO in DNA Replication and Replication Stress. Front Genet 7, 87 (2016).

3. W. Leung, R. M. Baxley, G. L. Moldovan, A. K. Bielinsky, Mechanisms of DNA Damage Tolerance: Post-Translational Regulation of PCNA. Genes (Basel) 10, (2018).

4. S. P. Bell, K. Labib, Chromosome Duplication in Saccharomyces cerevisiae. Genetics 203, 1027–1067 (2016).

5. P. M. J. Burgers, T. A. Kunkel, Eukaryotic DNA Replication Fork. Annu Rev Biochem 86, 417–438 (2017).

6. Y. Daigaku, A. A. Davies, H. D. Ulrich, Ubiquitin-dependent DNA damage bypass is separable from genome replication. Nature 465, 951–955 (2010).

7. M. K. Zeman, K. A. Cimprich, Causes and consequences of replication stress. Nat Cell Biol 16, 2–9 (2014).

8. A. A. Davies, D. Huttner, Y. Daigaku, S. Chen, H. D. Ulrich, Activation of ubiquitin-dependent DNA damage bypass is mediated by replication protein a. Mol Cell 29, 625–636 (2008).

9. C. Hoege, B. Pfander, G. L. Moldovan, G. Pyrowolakis, S. Jentsch, RAD6-dependent DNA repair is linked to modification of PCNA by ubiquitin and SUMO. Nature 419, 135–141 (2002).

10. P. Stelter, H. D. Ulrich, Control of spontaneous and damage-induced mutagenesis by SUMO and ubiquitin conjugation. Nature 425, 188–191 (2003).

11. J. E. Sale, Translesion DNA synthesis and mutagenesis in eukaryotes. Cold Spring Harb Perspect Biol 5, a012708 (2013).

12. D. Branzei, F. Vanoli, M. Foiani, SUMOylation regulates Rad18-mediated template switch. Nature 456, 915–920 (2008).

13. E. C. Minca, D. Kowalski, Multiple Rad5 activities mediate sister chromatid recombination to bypass DNA damage at stalled replication forks. Mol Cell 38, 649–661 (2010).

14. D. Gallo et al., Rad5 Recruits Error-Prone DNA Polymerases for Mutagenic Repair of ssDNA Gaps on Undamaged Templates. Mol Cell 73, 900–914 e909 (2019).

15. L. Kuang et al., A non-catalytic function of Rev1 in translesion DNA synthesis and mutagenesis is mediated by its stable interaction with Rad5. DNA Repair (Amst) 12, 27–37 (2013).

16. V. Pages et al., Requirement of Rad5 for DNA polymerase zeta-dependent translesion synthesis in Saccharomyces cerevisiae. Genetics 180, 73–82 (2008).

17. D. Gallo, G. W. Brown, Post-replication repair: Rad5/HLTF regulation, activity on undamaged templates, and relationship to cancer. Crit Rev Biochem Mol Biol 54, 301–332 (2019).

18. V. Alvarez et al., Orderly progression through S-phase requires dynamic ubiquitylation and deubiquitylation of PCNA. Sci Rep 6, 25513 (2016).

19. A. Gallego-Sanchez, S. Andres, F. Conde, P. A. San-Segundo, A. Bueno, Reversal of PCNA ubiquitylation by Ubp10 in Saccharomyces cerevisiae. PLoS Genet 8, e1002826 (2012).

20. V. Alvarez et al., PCNA Deubiquitylases Control DNA Damage Bypass at Replication Forks. Cell Rep 29, 1323–1335 e1325 (2019).

21. T. Kubota, Y. Katou, R. Nakato, K. Shirahige, A. D. Donaldson, Replication-Coupled PCNA Unloading by the Elg1 Complex Occurs Genome-wide and Requires Okazaki Fragment Ligation. Cell Rep 12, 774–787 (2015).

22. T. Kubota, K. Nishimura, M. T. Kanemaki, A. D. Donaldson, The Elg1 replication factor C-like complex functions in PCNA unloading during DNA replication. Mol Cell 50, 273–280 (2013).

23. E. Papouli et al., Crosstalk between SUMO and ubiquitin on PCNA is mediated by recruitment of the helicase Srs2p. Mol Cell 19, 123–133 (2005).

24. B. Pfander, G. L. Moldovan, M. Sacher, C. Hoege, S. Jentsch, SUMO-modified PCNA recruits Srs2 to prevent recombination during S phase. Nature 436, 428–433 (2005).

25. C. Yu et al., Strand-specific analysis shows protein binding at replication forks and PCNA unloading from lagging strands when forks stall. Mol Cell 56, 551–563 (2014).

26. T. A. Guilliam, J. T. P. Yeeles, An updated perspective on the polymerase division of labor during eukaryotic DNA replication. Crit Rev Biochem Mol Biol 55, 469–481 (2020).

27. S. A. Lujan, J. S. Williams, T. A. Kunkel, DNA Polymerases Divide the Labor of Genome Replication. Trends Cell Biol 26, 640–654 (2016).

28. P. Sung, Catalysis of ATP-dependent homologous DNA pairing and strand exchange by yeast RAD51 protein. Science 265, 1241–1243 (1994).

29. J. Zamarreño et al., Timely lagging strand maturation relies on Up-mediated PCNA dissociation from replicating chromatin. bioRxiv, (2024).

30. T. A. Guilliam, J. T. P. Yeeles, Reconstitution of translesion synthesis reveals a mechanism of eukaryotic DNA replication restart. Nat Struct Mol Biol 27, 450–460 (2020).

31. R. P. Wong, N. Garcia-Rodriguez, N. Zilio, M. Hanulova, H. D. Ulrich, Processing of DNA Polymerase-Blocking Lesions during Genome Replication Is Spatially and Temporally Segregated from Replication Forks. Mol Cell 77, 3–16 e14 (2020).

32. M. R. G. Taylor, J. T. P. Yeeles, The Initial Response of a Eukaryotic Replisome to DNA Damage. Mol Cell 70, 1067–1080 e1012 (2018).

33. Z. X. Zhou, S. A. Lujan, A. B. Burkholder, M. A. Garbacz, T. A. Kunkel, Roles for DNA polymerase delta in initiating and terminating leading strand DNA replication. Nat Commun 10, 3992 (2019).

34. A. Quinet, S. Tirman, E. Cybulla, A. Meroni, A. Vindigni, To skip or not to skip: choosing repriming to tolerate DNA damage. Mol Cell 81, 649–658 (2021).

35. C. Mellor, J. Nassar, S. Svikovic, J. E. Sale, PRIMPOL ensures robust handoff between on-the-fly and post-replicative DNA lesion bypass. Nucleic Acids Res, (2023).

36. M. S. Longtine et al., Additional modules for versatile and economical PCR-based gene deletion and modification in Saccharomyces cerevisiae. Yeast 14, 953–961 (1998).

37. R. Bermejo et al., Top1- and Top2-mediated topological transitions at replication forks ensure fork progression and stability and prevent DNA damage checkpoint activation. Genes Dev 21, 1921–1936 (2007).

38. C. Yu, H. Gan, Z. Zhang, Strand-Specific Analysis of DNA Synthesis and Proteins Association with DNA Replication Forks in Budding Yeast. Methods Mol Biol 1672, 227–238 (2018).

39. Y. Katou et al., S-phase checkpoint proteins Tof1 and Mrc1 form a stable replication-pausing complex. Nature 424, 1078–1083 (2003).

40. R. Bermejo, Y. M. Katou, K. Shirahige, M. Foiani, ChIP-on-chip analysis of DNA topoisomerases. Methods Mol Biol 582, 103–118 (2009).

41. B. Langmead, S. L. Salzberg, Fast gapped-read alignment with Bowtie 2. Nat Methods 9, 357–359 (2012).

42. H. Li et al., The Sequence Alignment/Map format and SAMtools. Bioinformatics 25, 2078–2079 (2009).

43. A. R. Quinlan, I. M. Hall, BEDTools: a flexible suite of utilities for comparing genomic features. Bioinformatics 26, 841–842 (2010).

44. C. C. Siow, S. R. Nieduszynska, C. A. Muller, C. A. Nieduszynski, OriDB, the DNA replication origin database updated and extended. Nucleic Acids Res 40, D682–686 (2012).

45. Y. Zhang et al., Model-based analysis of ChIP-Seq (MACS). Genome Biol 9, R137 (2008).

46. R-Core-Team, R: A language and environment for statistical computing., (2021).

47. M. Al Mamun, R. Bermejo. (https://github.com/MohammedAlMamun/ngsAnalyser1.1.4, 2022).

48. A. De Antoni, D. Gallwitz, A novel multi-purpose cassette for repeated integrative epitope tagging of genes in Saccharomyces cerevisiae. Gene 246, 179–185 (2000).

