## Supplemental Material for "Strand asymmetry of DNA damage tolerance mechanisms"

**The PDF file includes:**

Materials and Methods

Tables S1 and S2

Figs. S1 to S4

Materials and Methods

**Yeast strains and growth conditions.**

All strains are *RAD5^+^* derivatives of W303 and are listed in Table S1. Tagged alleles and gene deletions were generated by the single step PCR-based gene modification strategy **(*36*)**, by using plasmids and oligos listed in Table S2 and Table S3, respectively. The resulting genome constructions were checked by PCR. Additionally, protein tagging was also confirmed by trichloroacetic acid precipitation and western blot **(*37*)**.

Yeast cells were grown up to 1x10^7^ cells/ml in 200 mL of SD -URA at 23ºC and shifted to YPDA medium with 5 μg/mL α factor (Insight Biotechnology). G1 synchronization was monitored under the microscope (Motic) and cells were released into fresh YPDA medium supplemented with 0.2 mM HU (Ibian Technologies, REF HDU0250) or 0,033% MMS (Sigma, REF 129925) and 200 μg/mL BrdU (Sigma-Aldrich, REF B5002) for the times indicated. BrdU was not added in qPCR-ChIP experiments.

**eSPAN (*enrichment and Sequencing of Protein-Associated Nascent DNA*).**

eSPANs experiments were performed as described, with some modifications **(*38-40*)**. Cells were fixed with 1% formaldehyde for 30 minutes at room temperature, washed trice in 1X TBS, resuspended in lysis buffer (50 mM HEPES KOH pH 7,5, 140 mM NaCl, 1 mM EDTA, 1% Triton-X100, 0,1% sodium deoxycholate, 1mM PMSF, 1X antiproteolytic cocktail) and lysed with glass beads. Chromatin was then sonicated to an average DNA fragment size around 300 bp in length and immunoprecipitated using specific antibodies (listed in Table S4) attached to magnetic beads. Prior to immunoprecipitation, one fifth of chromatin was kept to process input and BrdU fractions. Immunoprecipitated chromatin was washed twice in lysis buffer, twice in lysis buffer supplemented with 360 mM NaCl, twice in wash buffer (10 mM Tris-HCl pH 8, 250 mM LiCl, 1 mM EDTA, 0,5 % NP-40, 0.5 % sodium deoxycholate) and once in TE 1X pH 8. Chromatin was then eluted from the magnetic beads in elution buffer (10 mM Tris-HCL pH 8, 10 mM EDTA, 1% SDS) and de-crosslinked overnight at 65 ºC. Samples were treated with 0,5 mg/mL proteinase K and incubated at 37 ºC for 2 hours followed by two phenol-chloroform purifications and precipitated with ethanol. Pellet was resuspended in 1X TE with 0,3 mg/mL RNase A, incubated at 37 ºC for 1 hour and purified with QIAquick PCR Purification Kit (Quiagen, REF 28104) following the manufacturer’s instructions. Eluted DNA and BrdU fraction were denatured at 100 ºC for 10 minutes and rapidly chilled on ice, followed by immunoprecipitation with anti-BrdU antibody attached to magnetic beads overnight. Nascent DNA was then washed and eluted in 1X TE as described in the first immunoprecipitation and incubated with 0,5 mg/mL proteinase K at 37 ºC for 1 hour. Samples were finally purified with QIAquick PCR Purification Kit. Prior to the second immunoprecipitation, 20 µL of sample were kept to amplify input and ChIP fractions.

**ssDNA library preparation and sequencing**

Libraries for sequencing were amplified from the different fractions using xGen™ ssDNA & Low-Input DNA Library Preparation Kit (Integrated DNA Technologies) following the manufacturer’s instructions, purified by size selection using AMPure XP beads (Beckman Coulter) and sequenced on Illumina NextSeq2000 platform using a paired-end sequencing strategy.

**Bioinformatics analysis.**

Raw sequencing reads for Input, ChIP, BrDU and eSPAN libraries were first aligned to reference yeast genome (S288C) using Bowtie2 version 2.4.4 **(*41*)**. Only properly aligned read pairs passing the mapping quality threshold (>= 30) were considered for further analysis. Resulting bam files were filtered with Samtools’ view, and bedtools *genomecoverage* function was used to calculate the strand-wise per base read coverage genome-wide **(*42, 43*)**. We used a sliding window of 10 base-pairs to bin the genome into windows spanning 300 bps (the approximate mean DNA fragment size in the experiment) and used bedtools’ map function to calculate strand-wise binned read coverages which were used for secondary analysis. Library size normalisation was performed for ChIP and BrDU against corresponding Input and for eSPAN against the respective BrDU data. Thereafter ChIP and BrDU were normalised by Input and, similarly, eSPAN was normalised by BrDU in strand-wise fashion. Fired replication origins in each respective experiment were inferred from the set of significant peaks (p-value cut-off 10^-5^), determined by macs2 *callpeak* function, that overlapped known origin locations in the corresponding BrDU library against Input as control **(*44, 45*)**. Custom R based functions were used to calculate average profiles around activated origins for normalised (Watson/Crick) read enrichments and stranded ratios **(*46*)**. To determine lagging to leading bias for eSPAN at individual origins: each origin environment was divided into four quadrants Watson left (WL), Watson right (WR), Crick left (CL) and Crick right (CR) centring the midpoint of the ARS element. WL and CR forms the lagging strand synthesis compartment, while WR and CL corresponds to the leading strand synthesis **(*25*)**. Thus, lagging over leading ratios were calculated, for the log2 transformed ratios – value greater than 0 indicates lagging bias for the given origin while value below 0 suggests leading bias. Prior to assigning a lagging or leading bias, we compared (WL + CR) to (WR + CL) and only if the binomial p-value was less than 10^-5^ – a bias pattern was assigned, otherwise the origin was defined to be indeterminate. Custom plotting functions in R base were used to draw the stranded read coverage at chromosome segments, average read enrichments and stranded ratios around active origins, and dot plots showing bias for all active origins. Except for coverage profile across given chromosomal coordinates, all the rest of analysis from sequence alignments to plotting were automated with the help of in-house software ngsAnalyser-1.1.4 **(*47*)**.

**Quantitative-PCR ChIP**

ChIP were performed as described in **(*39*)** and for the first immunoprecipitation of eSPAN experiment. Briefly, cells were fixed in 1 % formaldehyde for 30 minutes at room temperature and disrupted with glass beads. Chromatin was sonicated to an average DNA fragment size around 300 bp in length and immunoprecipitated using specific antibodies (listed in Table S4) attached to magnetic beads. Immunoprecipitated chromatin was extensively washed, treated with proteinase K, purified with phenol-chloroform and precipitated with ethanol. Pellet was resuspended in 1X TE with 0,3 mg/mL RNase A and purified with QIAquick PCR Purification Kit following the manufacturer’s instructions. Prior to immunoprecipitation, 20 µL of chromatin was kept to obtain the input fraction. Real time PCR was performed following a standard protocol. Primers used (listed in Table S3) amplify a region close to *ARS305, ARS306, ARS607* early replication origins and a distal site 15 kb away from *ARS305*. Input was used for normalizing immunoprecipitated DNA. A non-replicated region 15 Kb away from *ARS305* was used to normalize the chipped signal at replicated regions (*ARS305, ARS306* and *ARS607*), except for Fig S4 (in which ARS305 +15 Kb results are plotted).

**Table S1.**

| **Strain** | **Number** | **Genotype** | **Reference** |
| --- | --- | --- | --- |
| WT | RB3450 | *Mata RAD5 ade2-1 can1-100 his3-11,15 leu2-3,112 trp1-1 ura3::URA3/GPD-TK(7x)* | Lab stock |
| *rad5∆* | RB4149 | *Mata RAD5 ade2-1 can1-100 his3-11,15 leu2-3,112 trp1-1 ura3::URA3/GPD-TK(7x) rad5::TRP1* | This study |
| *rad18∆* | RB4146 | *Mata RAD5 ade2-1 can1-100 his3-11,15 leu2-3,112 trp1-1 ura3::URA3/GPD-TK(7x) rad18::HIS3MX6* | This study |
| *rad5∆ srs2∆* | RB5143 | *Mata RAD5 ade2-1 can1-100 his3-11,15 leu2-3,112 trp1-1 ura3::URA3/GPD-TK(7x) rad5::TRP1 srs2::HIS3MX6* | This study |
| *rad18∆ srs2∆* | RB5145 | *Mata RAD5 ade2-1 can1-100 his3-11,15 leu2-3,112 trp1-1 ura3::URA3/GPD-TK(7x) rad18::HIS3MX6 srs2::HIS3MX6* | This study |
| *rev1-FLAG* | RB4195 | *Mata RAD5 ade2-1 can1-100 his3-11,15 leu2-3,112 trp1-1 ura3::URA3/GPD-TK(7x) rev1::REV1::10FLAG::KANMX6* | This study |
| *rev1-FLAG* *rad5∆* | RB4272 | *Mata RAD5 ade2-1 can1-100 his3-11,15 leu2-3,112 trp1-1 ura3::URA3/GPD-TK(7x) rev1::REV1::10FLAG::KANMX6 rad5::TRP1* | This study |
| *rev1-FLAG* *rad18∆* | RB4268 | *Mata RAD5 ade2-1 can1-100 his3-11,15 leu2-3,112 trp1-1 ura3::URA3/GPD-TK(7x) rev1::REV1::10FLAG::KANMX6 rad18::HIS3MX6* | This study |
| *pol30-FLAG* | RB4197 | *Mata RAD5 ade2-1 can1-100 his3-11,15 leu2-3,112 trp1-1 ura3::URA3/GPD-TK(7x) pol30::POL30::10FLAG::KANMX6* | This study |
| *rad5-FLAG* | RB3983 | *Mata RAD5 ade2-1 can1-100 his3-11,15 leu2-3,112 trp1-1 ura3::URA3/GPD-TK(7x) rad5::RAD5::10FLAG::KANMX6* | This study |
| *rad5-MYC* | RB3969 | *Mata RAD5 ade2-1 can1-100 his3-11,15 leu2-3,112 trp1-1 ura3::URA3/GPD-TK(7x) rad5::RAD5::* *13MYC-HIS3MX6* | This study |
| *rad18-FLAG* | RB3932 | *Mata RAD5 ade2-1 can1-100 his3-11,15 leu2-3,112 trp1-1 ura3::URA3/GPD-TK(7x) rad18::RAD18::10FLAG::KANMX6* | This study |
| *ubp10-MYC* | RB4402 | *Mata RAD5 ade2-1 can1-100 his3-11,15 leu2-3,112 trp1-1 ura3::URA3/GPD-TK(7x) ubp10::UBP10::13MYC::HPHMX4* | This study |
| *ubp12-MYC* | RB4716 | *Mata RAD5 ade2-1 can1-100 his3-11,15 leu2-3,112 trp1-1 ura3::URA3/GPD-TK(7x) ubp12::UBP12::13MYC:: HPHMX4* | This study |
| *ubp10,12∆* | RB5058 | *Mata RAD5 ade2-1 can1-100 his3-11,15 leu2-3,112 trp1-1 ura3::URA3/GPD-TK(7x) ubp10::NATMX6 ubp12::HPHMX4* | This study |
| *rad30-FLAG* | RB4820 | *Mata RAD5 ade2-1 can1-100 his3-11,15 leu2-3,112 trp1-1 ura3::URA3/GPD-TK(7x) rad30::RAD30::10FLAG::KANMX6* | This study |
| *pol2-PK* | RB2160 | *Mata RAD5 ADE2 CAN1 his3-11,15 leu2-3, 112 trp1-1 ura3::URA3/GPD-TK(7X) pol2::POL2::6PK-HIS3MX6* | This study |
| *pol3-PK* | RB2164 | *Mata RAD5 ADE2 CAN1 his3-11,15 leu2-3, 112 trp1-1 ura3::URA3/GPD-TK(7X) pol3::POL3::9PK-TRP1* | This study |
| *rfa1-PK* | RB5209 | *Mata RAD5 ade2-1 can1-100 his3-11,15 leu2-3,112 trp1-1 ura3::URA3/GPD-TK(7x) rfa1::RFA1::9PK::HIS3MX6* | This study |

**Table S2.**

| **Plasmid** | **Use** | **Source** |
| --- | --- | --- |
| pU6H3-*FLAG* | To tag proteins with 10xFLAG epitope in the C-terminal end | Modified from **(*48*)** |
| pFA6a-*13MYC*-*TRP1* | To tag proteins with MYC epitope in the C-terminal end | **(*36*)** |
| pFA6a-*KANMX6* | To delete genes with *KAN* gene cassette | **(*36*)** |
| pFA6a-*TRP1* | To delete genes with *TRP1* gene cassette | **(*36*)** |
| pFA6a-*HIS3MX6* | To delete genes with *HIS3* gene cassette | **(*36*)** |
| pAG25-*NATMX6* | To delete genes with *NAT* gene cassette | Lab collection |

**Table S3.**

| **Oligo** | **Sequence (5’🡪3’)** | **Use** |
| --- | --- | --- |
| Rad5D pFA6a F | TTGAACAGGAAGAAAGGAAGAGGTTTTTTAACGATGACCTCGGATCCCCGGGTTAATTAA | To delete *RAD5* gene |
| Rad5 pFA6a R | CTATTCAAACAGCATCTGGATTTCTTCAATTCTCCTTTTTGAATTCGAGCTCGTTTAAAC |  |
| Rad18 pFA6a F | ATGGACCACCAAATAACCACTGCAAGCGACTTCACGACTACGGATCCCCGGGTTAATTAA | To delete *RAD18* gene |
| Rad18 pFA6a R | TTAATTGTTACCGGGTGGGTCTTTACTATATTCATTCAAGGAATTCGAGCTCGTTTAAAC |  |
| Srs2D F | CATTCCAATTTGATCTTTCTTCTACCGGTACTTAGGGATAGCAACAGCTGAAGCTTCGTACGC | To delete *SRS2* gene |
| Srs2D R | AAACCGCCTCCAATAGTTGACGTAGTCAGGCATGAAAGTGCTACATAGGCCACTAGTGGATCTG |  |
| Rad5-FLAG F | AAGACGAGAGAAGAAAAAGGAGAATTGAAGAAATCCAGATGCTGTTTGAATCCCACCACCATCATCATCAC | To tag *RAD5* gene |
| Rad5-FLAG R | GGTTGAAAATAATAATAAATAAAGTCTTTATATATGAGTATGTGGTATGAACTATAGGGAGACCGGCAGATC |  |
| Rad18-FLAG F | GAGAATTAATGGACTTGAATGAATATAGTAAAGACCCACCCGGTAACAATTCCCACCACCATCATCATCAC | To tag *RAD18* gene |
| Rad18-FLAG R | TTAACAAATGTGCACAAGCTAACAAACAGGCCTGATTACATATACACACCACTATAGGGAGACCGGCAGATC |  |
| Rev1-FLAG F | TTACCAGACTGTGCGTAAACTTGACATGGACTTTGAAGTTTCCCACCACCATCATCATCAC | To tag *REV1* gene |
| Rev1-FLAG R | GCGTGTTTACTGTATGCTGAAATGTTTTTTTTTTTTTAATACTATAGGGAGACCGGCAGATC |  |
| Rad5-MYC F | AAGACGAGAGAAGAAAAAGGAGAATTGAAGAAATCCAGATGCTGTTTGAACGGATCCCCGGGTTAATTAA | To tag *RAD5* gene |
| Rad5-MYC R | GGTTGAAAATAATAATAAATAAAGTCTTTATATATGAGTATGTGGTATGACGAATTCGAGCTCGTTTAAAC |  |
| Pol30-FLAG F | CCTACAGTTTTTCTTGGCTCCTAAATTTAATGACGAAGAATCCCACCACCATCATCATCAC | To tag *POL30* gene |
| Pol30-FLAG R | TTTATTATTTTTAGTATACAACTATATAGATAATTTACATACTATAGGGAGACCGGCAGATC |  |
| Rad30-FLAG F | ATCTTCCAAAAACATCTTATCATTTTTTACAAGAAAAAAATCCCACCACCATCATCATCAC | To tag *RAD30* gene |
| Rad30-FLAG R | TTTAGTTGCTGAAGCCATATAATTGTCTATTTGGAATAGGACTATAGGGAGACCGGCAGATC |  |
| ARS305-FwII | GTAACTTACACGGGGGCTAA | qPCR |
| ARS305-RV-II | ACTTTGATGAGGTCTCTAGC | qPCR |
| ARS305+15 kb Fwd | CCACCGAAACTTTGTGGCAAG | qPCR |
| ARS 305+ 15 kb Rv | GCACATGAGTACCCTCAATT | qPCR |
| ARS306 Fwd | GGACAAGGTGCAAATGCCAAG | qPCR |
| ARS306 Rv | CCCGCTCCTTCTCCTAACAT | qPCR |
| ARS607 FWD | CACATTATTCGGCACAGTAGG | qPCR |
| ARS607 Rv | GTGTCGCAGTCCATAGAAGG | qPCR |

**Table S4.**

| **Antibody** | **Source** | **Reference** |
| --- | --- | --- |
| ANTI-FLAG® M2 monoclonal antibody | Sigma-Aldrich | F1804 |
| Anti-Bromodeoxyuridine mAb | Bionova | MI-11-3 |
| Anti-Myc Tag Antibody, clone 4A6 | MERCK | 05-724 |
| Anti PK anti-V5 Tag antibody | Bionova | MCA1360 |
| Anti-Rad51 | Gift from Patrick Sung | **(*28*)** |

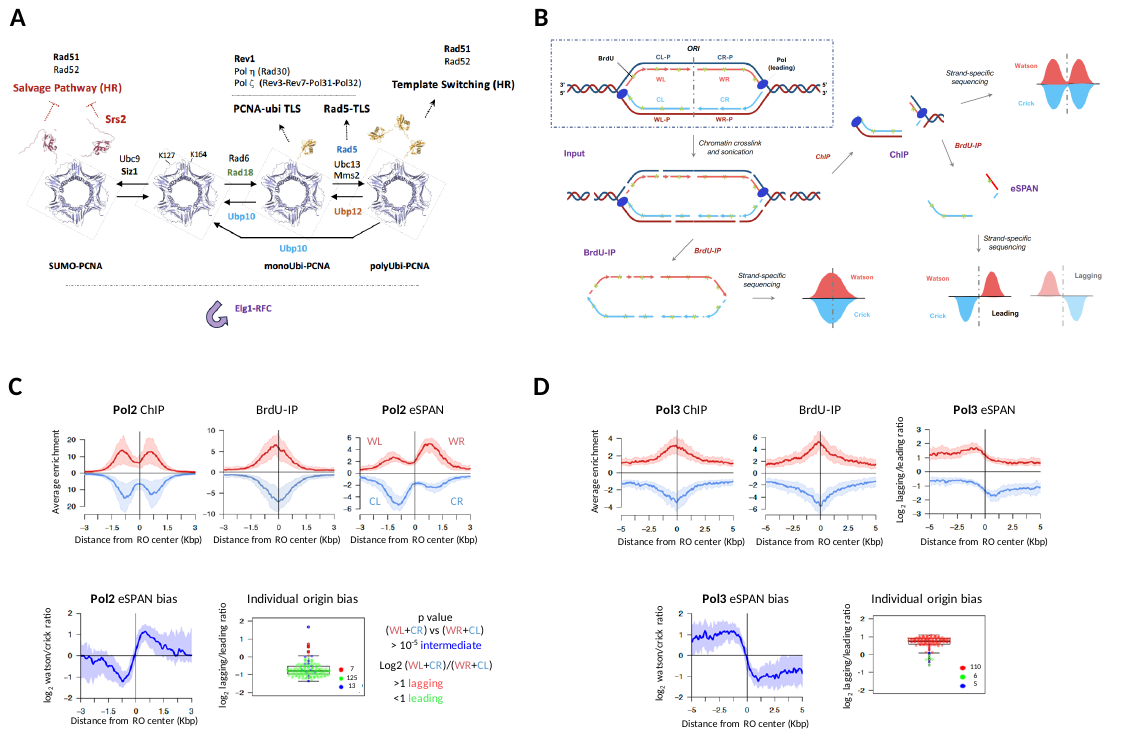

**Fig. S1. eSPAN for the analysis of DNA damage tolerance factor binding to stalled nascent DNA strands.** (**A**) Factors mediating PCNA post-translational modifications, DNA damage tolerance mechanisms and effectors (see text for details). (**B**) Schematic outline of the eSPAN methodology. Cells synchronously replicating incorporate BrdU into nascent chromatin, which is formaldehyde-crosslinked and sheared by sonication. BrdU-substituted DNA (BrdU-IP) and chromatin immunoprecipitation (ChIP) targeting a protein of interest (PoI) are then performed. Subsequently, an eSPAN (protein-associated nascent DNA) fraction is isolated by immunoprecipitation of BrdU-substituted DNA from the ChIP fraction. DNA from Input, BrdU, ChIP and eSPAN fractions is cleaned-up and subject to strand-specific sequencing. Reads corresponding to Watson (in red) and Crick (in blue) (P, parental) strands are mapped to the reference genome and aligned to active replication origins in order to plot enrichment profiles. Owing to the polarity of DNA polymerase synthesis, leading strand synthesis generates Crick DNA on leftwards moving forks (CL) and Watson DNA on rightwards moving forks (WR), while lagging strand synthesis produces Watson and Crick DNA on leftwards (WL) and rightward (CR) moving forks, respectively. Hence, lagging and leading nascent strand association to proteins of interest is estimated both globally and at individual replication origins. (**C-D**) eSPAN analysis of Pol2 (**C**) and Pol3 (**D**), catalytic subunits of DNA polymerase ε and δ, association to sites of nascent strand synthesis stalling. Cells were released from G1 in the presence of 0.2M HU and collected after one hour. Averaged input-normalized Pol2/Pol3-ChIP and BrdU-IP, as well as BrdU-IP-normalized Pol2/Pol3-eSPAN read enrichment ratios plotted around replication initiation sites. Median (red-blue) and 25-75 percentile (shaded) enrichment values plotted around BrdU-IP peak summits. Averaged Pol2/Pol3-eSPAN strand read ratios: median (deep blue) and 25-75 percentile (shaded) BrdU-IP-normalized Watson over Crick read log2 ratio values plotted around BrdU-IP peak summits. Individual replication origin leading-strand or lagging-strand biases: dot and box plots showing the variation of BrdU-IP normalized lagging over leading eSPAN read ratios around BrdU-IP peak summits. Each dot corresponds to an individual origin-containing eSPAN peak, falling into one of three categories (intermediate, lagging-strand bias or leading-strand bias) based on lagging over leading read binomial distribution significance and bias calculation (as shown on panel C).

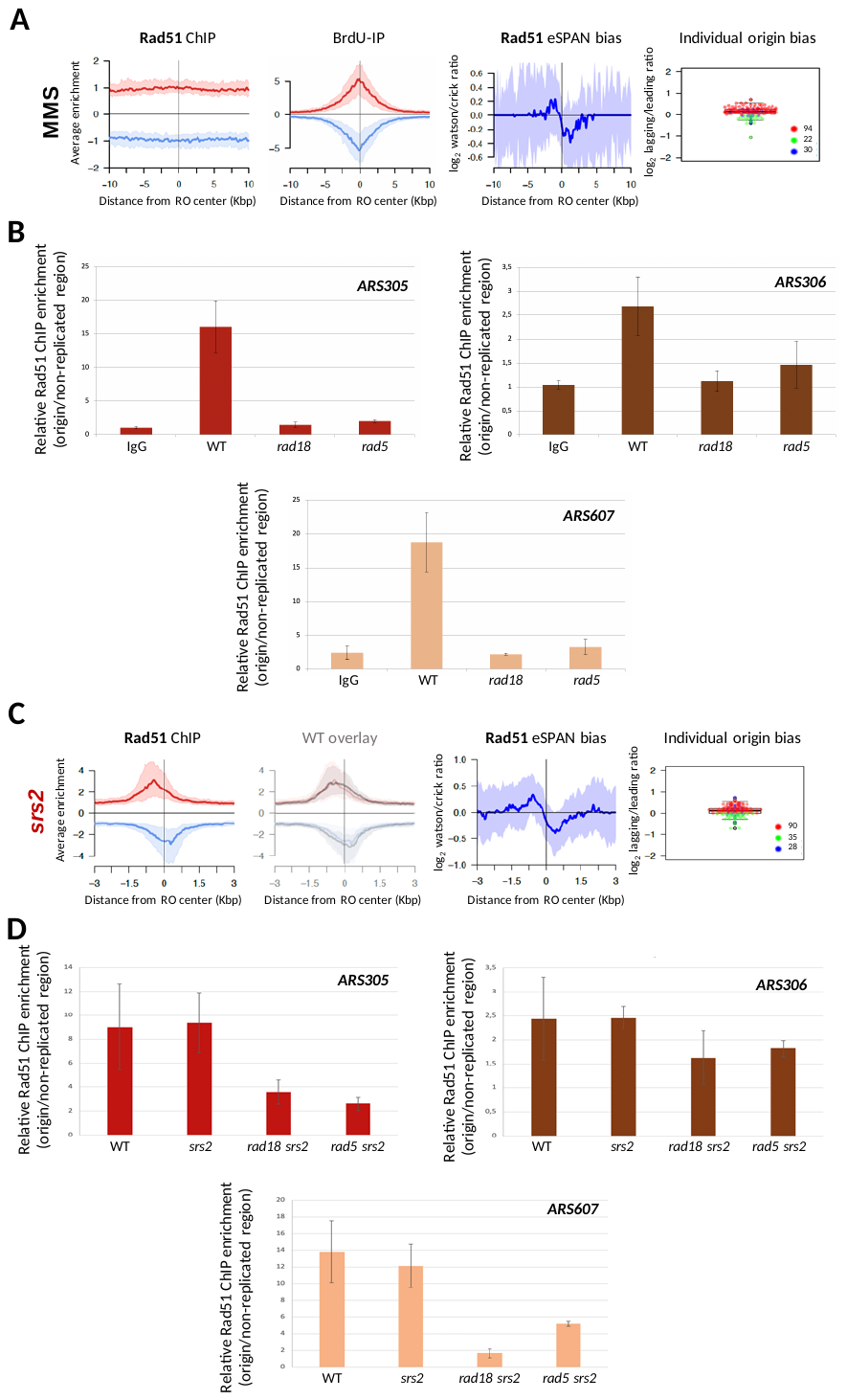

**Fig. S2. Genomic analysis of Rad51 association to sites of replicative polymerase stalling.** (**A**) Rad51-eSPAN analysis under MMS-induced polymerase stress. Cells were released from an α-factor-induced G1 block in the presence of 0,033% MMS and collected after 30 minutes. Averaged input-normalized Rad51-ChIP and BrdU-IP read enrichment profiles, as well as averaged Rad51-eSPAN strand read ratios around replication initiation sites and individual replication origin leading-strand or lagging-strand biases are shown. (**B**). Chromatin immunoprecipitation followed by quantitative PCR (ChIP-qPCR) analysis of Rad51 binding to sites of nascent strand synthesis stalling. Cells were released from G1 in the presence of 0.2M HU and collected after one hour for ChIP using anti-Rad51 or IgG control antibodies. Histogram plots show relative Rad51 enrichment at sites of polymerase stalling close to the early replication origins *ARS305*, *ARS306* and *ARS607* normalized against a region (*ARS305 +15Kb*) that is not replicated in these conditions. Mean and SD values from three independent experiments are shown. (**C**) eSPAN analysis of Rad51 association to replicative polymerase stalling sites in *srs2Δ* cells corresponding to the experiment shown in Fig. 1F. (**D**) ChIP-qPCR analysis of Rad51 biding to sites of nascent strand synthesis stalling in WT, *srs2Δ*, *rad18Δ* *srs2Δ* and *rad15Δ* *srs2Δ* cells as in B. Mean and SD values from three independent experiments are shown.

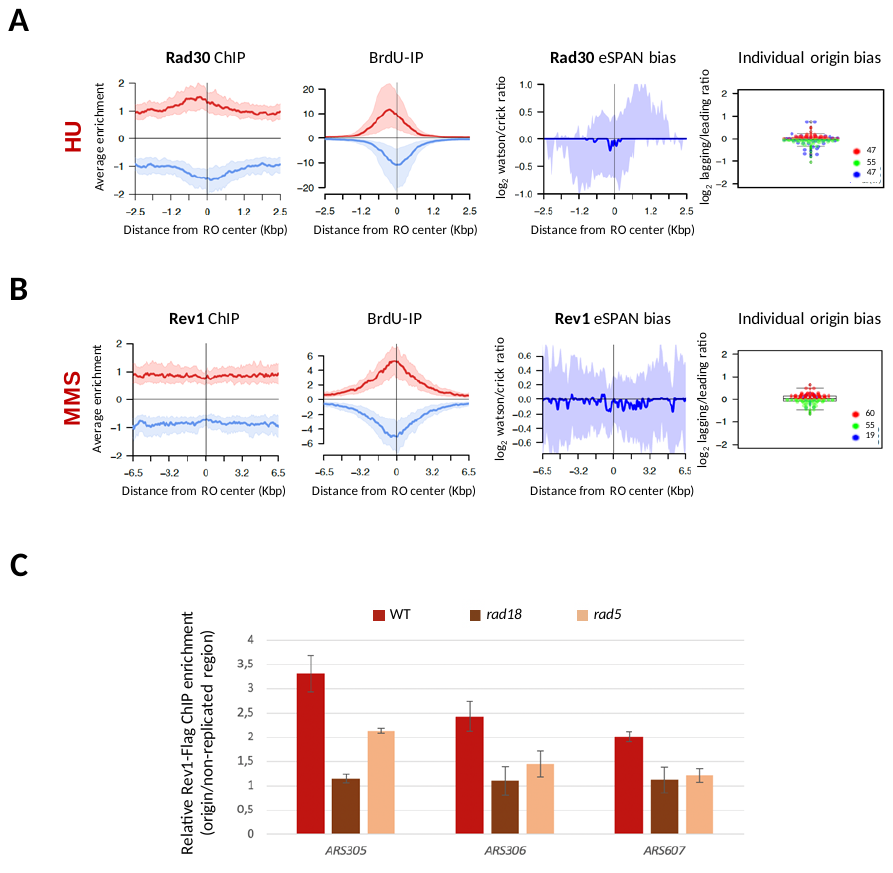

**Fig. S3. Genomic analysis of TLS polymerase association to sites of replicative polymerase stalling.** (**A**) eSPAN analysis of DNA polymerase η association to replicative polymerase stalling sites. Rad30-Flag cells were released from G1 in the presence of 0.2M HU and collected after one hour. Averaged input-normalized Rad30-ChIP and BrdU-IP read enrichment profiles, as well as averaged Rad30-eSPAN strand read ratios and individual replication origin leading-strand or lagging-strand biases are shown. (**B**) eSPAN analysis of Rev1 under MMS-induced polymerase stress. Cells were released from G1 block in the presence of 0.033% MMS and collected after 30 minutes. Averaged input-normalized Rev1-Flag-ChIP and BrdU-IP read enrichment profiles, as well as averaged Rev1-Flag-eSPAN strand read ratios and individual replication origin leading-strand or lagging-strand biases are shown. (**C**) Chromatin immunoprecipitation followed by quantitative PCR (ChIP-qPCR) analysis of Rev1 biding to sites of nascent strand synthesis stalling in Rev1-Flag WT, *rad18Δ* and *rad5Δ* cells.

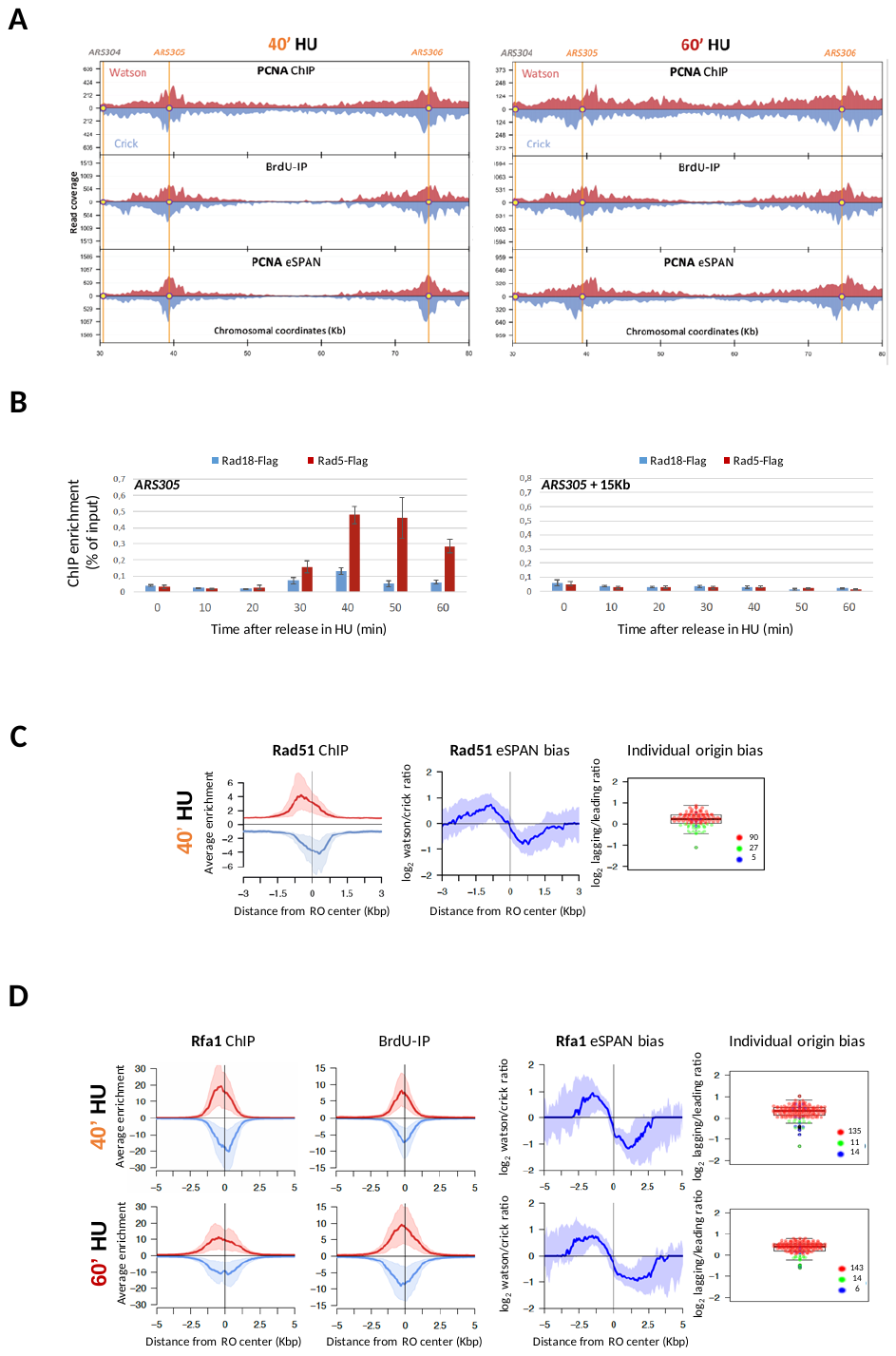

**Fig. S4. Genomic analysis of PCNA and PCNA writers to sites of replicative polymerase stalling.** (**A**) eSPAN analysis of PCNA as in Fig. 3A. PCNA-ChIP, BrdU-IP and PCNA-eSPAN fraction read coverage plotted along a region spanning from 30 to 80 Kb on chromosome III. (**B**) Analysis of Rad18 and Rad5 PCNA-K164 writer dynamic recruitment to sites of replicative polymerase stalling. ChIP-qPCR of Rad18-Flag and Rad5-Flag binding close to *ARS305* (*ARS305*) or at a 15Kb-distant unreplicated region (*ARS305* +15Kb) in cells released from G1 into 0.2M HU for the indicated times. Mean and SEM values from at least three independent experiments are shown. (**C**) Rad51-eSPAN analysis in cells collected 40 minutes after release from G1 in the presence of 0.2M HU. Averaged input-normalized Rad51-ChIP read enrichment profiles, as well as averaged Rad51-eSPAN strand read ratios and individual replication origin leading-strand or lagging-strand biases are shown. (**D**) eSPAN analysis of RPA association to replicative polymerase stalling sites. The large subunit of the RPA heterotrimeric complex Rfa1 was tested. Rfa1-PK cells were released from G1 in the presence of 0.2M HU and collected after 40 or 60 minutes. Averaged input-normalized Rfa1-ChIP and BrdU-IP read enrichment profiles, as well as averaged Rfa1-eSPAN strand read ratios and individual replication origin leading-strand or lagging-strand biases are shown.
